# In-Search Assignment of Monoisotopic Peaks Improves the Identification of Cross-Linked Peptides

**DOI:** 10.1101/335851

**Authors:** Swantje Lenz, Sven H. Giese, Lutz Fischer, Juri Rappsilber

## Abstract

Cross-linking/mass spectrometry (CLMS) has undergone a maturation process akin to standard proteomics by adapting key methods such as false discovery rate control and quantification. A seldom-used search setting in proteomics is the consideration of multiple (lighter) alternative values for the monoisotopic precursor mass to compensate for possible misassignments of the monoisotopic peak. Here, we show that monoisotopic peak assignment is a major weakness of current data handling approaches in cross-linking. Cross-linked peptides often have high precursor masses, which reduces the presence of the monoisotopic peak in the isotope envelope. Paired with generally low peak intensity, this generates a challenge that may not be completely solvable by precursor mass assignment routines. We therefore took an alternative route by ‘in-search assignment of the monoisotopic peak’ in Xi (Xi-MPA), which considers multiple precursor masses during database search. We compare and evaluate the performance of established preprocessing workflows that partly correct the monoisotopic peak and Xi-MPA on three publicly available datasets. Xi-MPA always delivered the highest number of identifications with ~2 to 4-fold increase of PSMs without compromising identification accuracy as determined by FDR estimation and comparison to crystallographic models.

---

Several approaches have been utilized to increase the numbers of identified cross-links, for example enriching for cross-linked peptides^1–3^, using different proteases^1–3^ or optimizing fragmentation methods^4,5^. In parallel with experimental developments, data analysis has also progressed. Search software has been designed for the identification of cross-linked peptides, for example Kojak^6^, xQuest^7^, pLink^8^, XlinkX^9^ or Xi^3^. In addition, cross-linking workflows can make use of preprocessing methods to improve data quality and reduce file sizes^10^, as well as post-processing methods to filter out false identifications^6,11^ and custom-tailored false discovery rate (FDR) estimation^12–14^. Preprocessing can improve peptide identification by correcting the MS1 precursor ion m/z and simplifying MS2 fragment spectra. Established proteomics software perform such preprocessing, including MaxQuant^15,16^ and OpenMS^17,18^. For example, MaxQuant performs a variety of preprocessing steps: it corrects the precursor m/z by an intensity-weighted average if a suitable peptide feature is found, reassigns the monoisotopic peak and contains options for intensity filtering of MS2 peaks. Despite such correction of the precursor mass, many linear search engines have integrated the possibility of considering multiple monoisotopic peaks during search^19–21^. However, the benefits of this search feature are currently unclear, and it is not in common use. It seems that the assignment of monoisotopic mass for tryptic peptides is already achieved adequately either during acquisition or as part of preprocessing.

Cross-linked peptides have characteristics that may render MS1 monoisotopic precursor mass assignment as used for linear peptides nonoptimal: high-charge states, large masses, and low abundances. Several cross-link search engines include MS1 correction in their pipeline: pLink^8^ corrects monoisotopic peaks based on previous work with linear peptides^22^. Kojak averages precursor ion signals of neighboring scans to create a composite spectrum and infer the true monoisotopic mass of the precursor. If this step fails, precursor masses up to −2 Da lighter are searched, albeit with currently unclear benefits. For previous searches in Xi, MaxQuant was used to perform preprocessing. Nevertheless, we are not aware of a detailed evaluation of the impact of different preprocessing techniques for cross-link identification. Correcting the monoisotopic mass of precursors, although acknowledged as an issue^6,23^, awaits systematic evaluation.

In this study, we show that errors in assigning monoisotopic peaks during data acquisition are frequent for cross-linked peptides because of their size and generally low abundance. This adversely affects their identification. We show that standard software suites, MaxQuant and OpenMS correct monoisotopic precursor masses of cross-linked peptides with variable success. We then implement an option in Xi to consider multiple precursor masses during search, in order to minimize the impact of false monoisotopic precursor mass assignment on the identification of cross-links.

## METHODS

### Datasets

In this study, we used three publicly available datasets (Table 1). The three datasets were chosen to reflect a range of applications of cross-linking mass spectrometry as well as a range of data complexity: the first dataset is Human Serum Albumin (HSA) cross-linked with succinimidyl 4,4-azipentanoate (SDA) and fragmented using five different methods (PXD003737)^24^. The second dataset is a pooled pseudo-complex sample with seven separately cross-linked proteins with bis(sulfosuccinimidyl) suberate (BS3) (PXD006131)^4^. This dataset includes data from four different fragmentation methods. The third dataset is the most complex sample, composed of 15 size exclusion chromatography fractions of *Chaetomium thermophilum* lysate cross-linked with BS3 and fragmented only with HCD (PXD006626)^25^. The first and last size exclusion fractions were used to optimize the search parameters for this dataset. All samples were analyzed on an Orbitrap Fusion Lumos Tribrid mass spectrometer (Thermo Fisher Scientific, San Jose, CA) using Xcalibur (version 2.0 and 2.1).

**Table 1.**
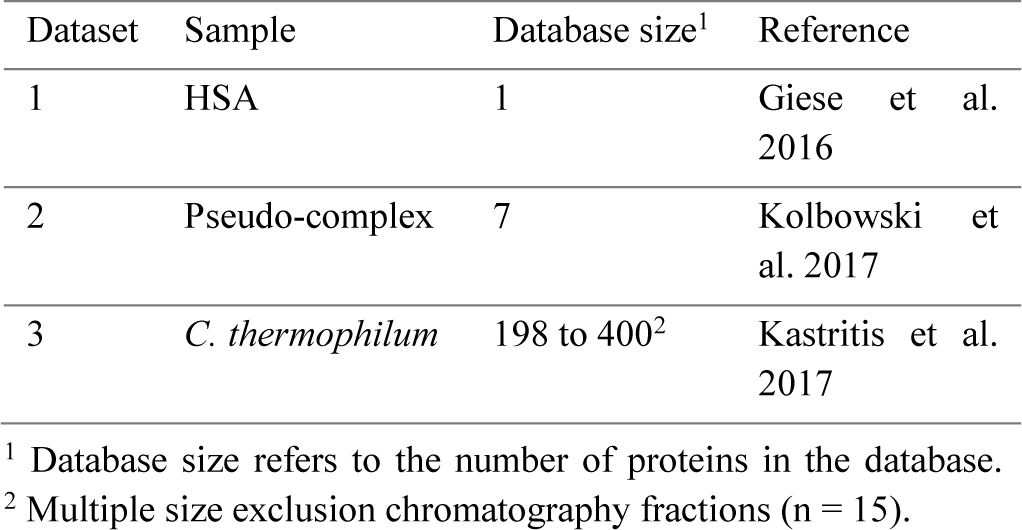
Overview of datasets used.

| Dataset | Sample | Database size<sup>1</sup> | Reference |
| --- | --- | --- | --- |
| 1 | HSA | 1 | Giese et al. 2016 |
| 2 | Pseudo-complex | 7 | Kolbowski et al. 2017 |
| 3 | *C. thermophilum* | 198 to 400<sup>2</sup> | Kastritis et al. 2017 |

<sup>1</sup> Database size refers to the number of proteins in the database.
<sup>2</sup> Multiple size exclusion chromatography fractions (n = 15).

### Preprocessing

Raw files were preprocessed independently using MaxQuant (1.5.5.30), OpenMS (2.0.1) and the ProteoWizard^26^ tool msconvert (3.0.9576) for comparison. Scripts automating the preprocessing, search and evaluation were written in Python (2.7).

The essential steps during the preprocessing can be divided into two parts: (1) correction of the m/z or charge of the precursor peak for MS2 spectra and (2) denoising of MS2 spectra. MaxQuant and OpenMS both try to correct the precursor information via additional feature finding steps, i.e. identifying a peptide feature from the retention time, m/z and intensity domain of the LC-MS run. Additionally, denoising of the MS2 spectra is performed by simply filtering the most intense peaks in defined m/z windows. The preprocessing is by default enabled in MaxQuant and was run using the partial processing option (steps 1 to 5) with default settings except for inactivated ‘deisotoping’ and ‘top peaks per 100 Da’, which was set to 20. The OpenMS preprocessing workflow includes centroiding, feature finding^27^, precursor correction (mass and charge) using the identified features and MS2 denoising as described above (**Figure S1**). Msconvert was used to convert the raw files to mgf files without any correction. These peak files were denoted as ‘uncorrected’ and used as our baseline to quantify improvements in the subsequent database search. For the ‘in-search assignment of the monoisotopic peak’ in Xi (Xi-MPA), we used msconvert to convert raw files to mgf files and included a MS2 peak filter for the 20 most intense peaks in a 100 m/z window.

### Data Analysis

Peak files were searched separately in Xi (1.6.731) with the following settings: MS accuracy 3 ppm, MS/MS accuracy 10 ppm, oxidation of methionine as variable modification, tryptic digestion, 2 missed cleavages. For samples cross-linked with SDA, linkage sites were allowed on lysine, serine, tyrosine, threonine and protein n-terminus on one end and all amino acids on the other end of the cross-linker. Variable modifications were mono-link SDA (110.048 Da), SDA loop-links (82.0419 Da), SDA hydrolyzed (100.0524 Da), SDA oxidized (98.0368 Da)^24^ as well as carbamidomethylation on cysteine. For searches with BS3, linkage sites were lysine, serine, threonine, tyrosine and the protein n-terminus. Carbamidomethylation on cysteine was set as fixed modification. Allowed variable modifications of the cross-linker were aminated BS3 (155.0946 Da), hydrolyzed BS3 (156.0786 Da) and loop-linked BS3 (138.0681 Da). For collision-induced dissociation (CID) and beam-type CID, also referred to as higher-energy C-trap dissociation (HCD), b- and y-ions were searched for, whereas for electron transfer dissociation (ETD) c- and z-ions were allowed. For ETciD and EThcD, b-, c-, z- and y-ions were allowed. The HSA and pseudo-complex datasets were searched against the known proteins in the sample. For each protein fraction of the *C. thermophilum* dataset, the databases of the original publication were used, where a database was created for each fraction by taking the most abundant proteins (iBAQ value above 10^6^). Datasets 1 and 2 were searched with a reversed decoy database, while dataset 3 was searched with a shuffled decoy database due to palindromic sequences.

For cross-linking, there are different information levels: PSMs, peptide pairs, residue pairs (links) and protein pairs. The false discovery rate (FDR) can be calculated on each one of these levels and should be reported for the level at which the information is given^12^. The FDR was calculated with xiFDR (1.0.14.34) and a 5% PSM level cutoff was imposed. The setting ‘uniquePSMs’ was enabled and the FDR was calculated separately on self and between links. Minimal peptide length was set to 6. In dataset 2, identified cross-linked residues were mapped to the crystal structure of the respective protein and the Euclidian distance between the alpha-carbons was calculated. Structures were downloaded from the PDB (IDs: 1AO6, 5GKN, 2CRK, 3NBS, 1OVT, 2FRJ).

## RESULTS AND DISCUSSION

We evaluated the impact on cross-link identification in Xi of changing the precursor monoisotopic mass that was initially assigned during data acquisition (‘uncorrected’). In this analysis, MaxQuant and OpenMS were used as preprocessing tools. We used three different datasets that differ in complexity and fragmentation regimes. To measure the improvements from using the preprocessing tools, a simple conversion from raw files to mgf format was done with msconvert and used as a baseline. Note that in the spectrum header, there are two m/z values: the trigger mass of the MS2 and the assigned monoisotopic peak of the isotope cluster. Msconvert extracts the assigned monoisotopic mass. Processed data were searched separately in Xi and evaluated on PSM or (unique residue pair) link level, with a 5% FDR. Finally, the newly implemented in-search assignment of monoisotopic peaks in Xi was compared to the elaborate preprocessing pipelines in OpenMS and MaxQuant.

### Preprocessing increases the number of cross-link PSMs by finding the correct monoisotopic peak

The datasets were preprocessed in MaxQuant and OpenMS and numbers of identified PSMs were compared to those obtained using uncorrected data. Datasets 1 (HSA) and 2 (pseudo-complex) were acquired with different acquisition methods. For comparability to dataset 3 (complex mixture), we focused on the HCD acquired data. Cross-links between proteins were excluded, either because they were experimentally not possible (dataset 2) or observed in too low numbers for reliable FDR calculation (dataset 3).

For uncorrected data, 672, 354, and 2157 cross-link PSMs resulted for the HSA dataset (dataset 1), pseudo-complex (dataset 2), and first and last fractions of *C. thermophilum* respectively (dataset 3). Both preprocessing approaches improved numbers of identified PSMs for all datasets: Preprocessing in MaxQuant led to 1127 (68% increase), 966 (173% increase) and 2966 (38% increase), while for OpenMS, 1044 (55% increase), 598 (69% increase) and 2394 (11% increase) PSMs were identified (Figure 1A).

**Figure 1.**
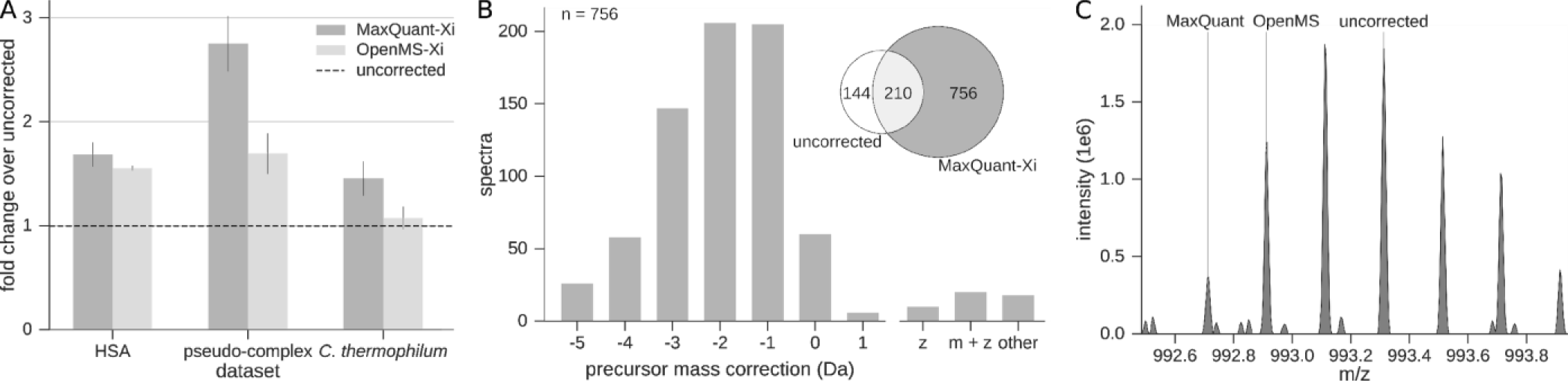
Correction of the monoisotopic peak is crucial in cross-link identification. (A) The datasets were preprocessed using MaxQuant and OpenMS, leading to more identified PSMs in all cases. Fold changes from uncorrected data (msconvert conversion of Xcalibur data) were calculated for each file separately and the mean plotted. Error bars represent the standard error of the mean between different acquisitions (HSA: n = 3, pseudo-complex: n = 3, *C. thermophilum*: n = 8). (B) The majority of additional identifications after preprocessing are due to correction of the precursor mass to lighter monoisotopic masses. Spectra that are unique to MaxQuant preprocessed searches of HCD acquisitions from dataset 2 were evaluated in terms of precursor correction. The main proportion of the gain was corrected to lighter masses of up to −3 Da, while charge state correction or correction to heavier masses rarely occurred. (C) Isotope cluster of a corrected precursor of m/z 992.71 (z = 5, m = 4958.6 Da) was solely identified in MaxQuant preprocessed results. In OpenMS preprocessed and uncorrected data, the wrong monoisotopic mass was selected for unknown reasons.

We assessed the gains in identified PSMs of preprocessed data compared to uncorrected data (focusing on dataset 2) regarding three forms of precursor correction: (1) correction of the monoisotopic mass, (2) charge state correction and (3) small corrections of the m/z value based on averaging the m/z values across the peptide feature (Figure 1B). Precursor mass and charge state of spectra identified solely in MaxQuant-Xi were compared to their counterparts when searching uncorrected data in Xi. Of the 756 newly identified spectra, 686 (91%) had a different monoisotopic precursor mass. Precursors were primarily corrected to lighter masses by MaxQuant, i.e. the monoisotopic peak correction by −1 (208 spectra), −2 (215 spectra), −3 (149 spectra) and −4 Da (62 spectra). Greater shifts (−5 to −7 Da) only occurred 30 times, and corrections to heavier masses were observed 22 times. Only 30 spectra (4%) were corrected in their charge state. For the 60 spectra (8%) without correction in charge state or monoisotopic peak, we only identified nine spectra that had a higher error than 3 ppm before preprocessing, indicating a small correction of the initial precursor m/z (by averaging of peptide feature peaks). The main proportion of these identifications is likely a product of noise removal in MS2 spectra or small changes in the score distribution. Similarly, for OpenMS-Xi, the monoisotopic peak correction had the greatest impact: Of the 314 spectra that OpenMS added over uncorrected data, 139 were precursor corrected by −1 Da and 108 to −2 Da. In contrast to MaxQuant, corrections to −3 or lighter were not observed, which might explain the higher number of identifications obtained with MaxQuant-Xi.

In summary, preprocessing, especially monoisotopic peak correction, leads to a notable increase in identifications. Using the 3-dimensional peptide feature is advantageous compared to on-the-fly detection of the monoisotopic peak. If the preceding MS1 spectrum was acquired during the beginning (or end) of the elution profile of a peptide, the intensity will be low. Thus, the monoisotopic peak might not even be detectable at the time of fragmentation. For large (cross-linked) peptides, this effect might be exacerbated by the monoisotopic peak usually being less intense than other isotope peaks. Therefore, using the additional information from the retention time domain will be beneficial. The same feature information can also be used to determine or validate the assigned charge state of the precursor. However, the instrument software almost always assigned the same charge state as MaxQuant or OpenMS. Thus, the main advantage for identifying cross-linked peptides arises from monoisotopic peak correction.

Interestingly, OpenMS and MaxQuant did not always agree on or find the same monoisotopic peak (Figure 1C). Of the total MaxQuant-corrected spectra with a different monoisotopic mass, 81% were not corrected and 6% corrected differently with OpenMS. Vice versa, 15% of the monoisotopic peaks corrected by OpenMS were not corrected by MaxQuant and 25% were corrected differently. Both MaxQuant and OpenMS have their own implementations for precursor correction - therefore, there might be instances where MaxQuant is able to find a corresponding peptide feature where OpenMS does not and vice versa. Although OpenMS did not lead to the same improvements in the number of identifications as MaxQuant, it did correct some precursors that the latter did not. We therefore suspect that there are also precursors with a falsely assigned monoisotopic peak that were corrected with neither algorithm. Furthermore, 3-dimensional detection of peptide features is challenging for low intensity peptides. In conclusion, there likely remain falsely assigned monoisotopic peaks in the data, ultimately leading to missed or false identifications.

### In-search monoisotopic peak assignment increases the number of identifications

We observed multiple cases where MaxQuant and OpenMS disagreed in their monoisotopic peak choice, indicating that the problem of monoisotopic peak assignment (MPA) cannot be solved easily at MS1 level. Indeed, we found instances where the monoisotopic peak is simply not distinguishable from noise, so a feature-based correction would not be feasible. Nevertheless, the associated MS2 spectra could be matched to a cross-linked peptide when considering multiple different monoisotopic masses during search. This shows that the extra information of obtaining a peptide-spectrum match is advantageous to MPA over considering MS1 information alone. Therefore, we implemented a monoisotopic peak assignment in Xi: for each MS2 spectrum, multiple precursor masses are considered during a single search and the highest scoring peptide-pair assigns the precursor mass. Note that this is different from simply searching with a wide mass error for MS1. The mass accuracy of MS1 is minimally compromised as multiple candidates for the monoisotopic mass are taken and considered with the original mass accuracy of the measurement.

To find a good trade-off between increased search space and sensitivity, we tested different mass range settings on the datasets. For dataset 2 (HCD subset), the number of PSMs increased with ranges up to −5 Da on the considered monoisotopic masses (Figure 2A). However, the increase in identifications from −4 to −5 Da was only 3% and considering the increase in search time, we continued with a maximal correction to −4 Da as the optimal setting for this dataset. Xi-MPA yielded 1508 PSMs, which is a 326% increase compared to searching uncorrected data and a 56% increase compared to MaxQuant-Xi. Similar improvements are observed for the other fragmentation methods in this dataset (**Figure S2**). Additionally, we corrected up to −7 Da to test if a large increase in search space increases random spectra matches as measured by the target-decoy approach. The number of identifications at 5% FDR decreased only slightly compared to −5 Da (−1%), but still led to more identifications than up to −4 Da (3%). In the HSA dataset, Xi-MPA with up to −4 Da increased the number of identified PSMs by 170% compared to uncorrected data (**Figure S3**).

**Figure 2.**
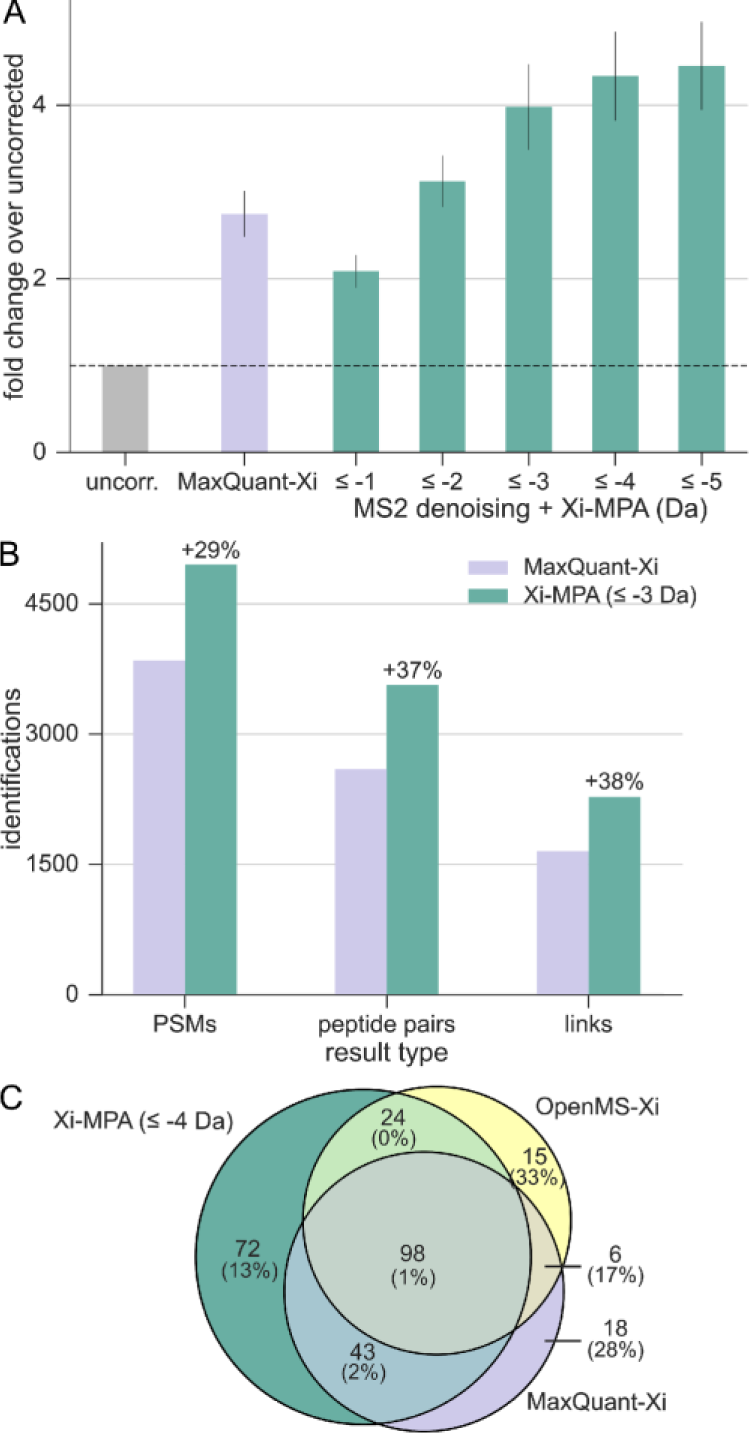
In-search monoisotopic peak assignment outperforms preprocessing. (A) Performance of Xi-MPA on dataset 2. HCD data from the pseudo-complex dataset were searched assuming different ranges of missing monoisotopic peaks. With increased ranges, the number of identified PSMs also increases. (B) Performance of Xi-MPA on the complete *C. thermophilum* dataset. All 15 fractions were searched with the original preprocessed data as well as with Xi-MPA. (C) Overlap of identified residue pairs of MaxQuant-Xi and OpenMS-Xi to residue pairs gained from Xi-MPA (dataset 2). Numbers in brackets are the proportion of decoys in the respective regions.

As a final evaluation of in-search monoisotopic peak assignment, we searched the complete dataset of *C. thermophilum*. We used 0 to −3 Da as the range of Xi-MPA, since an initial analysis of the first and last fraction of the *C. thermophilum* dataset returned a similar number of identifications when running Xi-MPA up to −4 Da or −3 Da (**Figure S4**). As a comparison, we took the original peak files obtained from PRIDE. The FDR was calculated separately on self and between links, with a minimum of 3 fragments per peptide and a minimal delta score of 0.5. For the original peak files, which were preprocessed in MaxQuant, we identified 3848 PSMs, 2594 peptide pairs and 1653 cross-links, with a 5% FDR on each respective level (Figure 2B). Xi-MPA resulted in 4952 PSMs (29% increase), 3566 peptide pairs (37% increase) and 2273 cross-links (38% increase).

Next, we looked into the complementarity of search results with the different approaches, using dataset 2 at 5% link-FDR. Preprocessing via MaxQuant and OpenMS led to 172 and 158 links, respectively, while Xi-MPA resulted in 243 links. While the overlap between links of OpenMS-Xi and MaxQuant-Xi is only 50%, Xi-MPA identifications cover 76% of both searches (Figure 2C). 19 and 23 links are uniquely found in MaxQuant and OpenMS preprocessed data respectively. However, there are 5 decoy links as well in each unique set (resulting in a link-FDR of 26% and 22%). For Xi-MPA, there are 75 unique target links with 12% link-FDR.

Identification-based monoisotopic peak assignment as employed by Xi-MPA results in more identifications than the feature-based assignment algorithms of OpenMS and MaxQuant. Neither OpenMS nor MaxQuant correct all precursor masses that are incorrectly assigned during data acquisition. In Xi-MPA, spectra are searched with multiple monoisotopic masses, thereby relying less on the MS1 information. The quality of the precursor isotope cluster does not contribute to the decision of monoisotopic mass and spectra for which correction failed will be identifiable. One could hypothesize that increasing the search space by considering multiple masses will lead to more false positives, thereby reducing the number of true identifications. This is not the case, as we match substantially more PSMs at constant FDR by considering alternative monoisotopic masses. As a second plausible caveat, this approach increases the search time. However, the use of relatively cheap computational time appears balanced by the notable increase in identified cross-links. The optimal range of additional monoisotopic peaks to search will however be dependent on complexity and quality of MS1 acquisition and the instrument software. To reduce the mass range considered in Xi-MPA, we developed a MS1 level-based approach. For each precursor, we search lighter isotope peaks in MS1 and use this to narrow the search space (explained in detail in Supporting Information). This led to an average of 24% less values to be considered, while only reducing the number of identifications by 3%. We hope that our observation of the monoisotopic peak detection challenge in cross-linking together with our publicly available datasets will lead to further improvements in monoisotopic peak-assignment algorithms in the future, possibly tailored to cross-link data.

### In-search monoisotopic peak assignment does not compromise search accuracy

Changing the search could lead to several problems. We already excluded that the increased search space leads to high-scoring decoy matches that in turn reduce the number of identifications at a given FDR cut-off. As an additional validation, we assessed our results against known PDB structures using the HCD data from the pseudo-complex dataset (dataset 2), at 5% link-FDR. Assuming a crystal structure is correct, a cross-link can be unexpectedly long either because the link is false or because of in-solution structural dynamics. If, however, the proportion of long-distance links in results of two approaches is identical, then at least the two results have equal quality.

We first tested the results of all three approaches against crystal structures. Residue pairs were mapped to PDB structures and the distance between the two alpha-carbons was calculated (see Methods). 30 Å was set as the maximal distance for BS3, links with a greater distance were classified as long-distance. In this evaluation, we excluded the protein C3B because its flexible regions make it unsuitable for this analysis. For MaxQuant and OpenMS preprocessed results, 11.8% and 6.1% long-distance cross-links were identified, respectively. In Xi-MPA, 8.1% long distance cross-links were identified (Figure 3A). Of the links uniquely identified through Xi-MPA, only 5.3% were long distance links. Therefore, Xi-MPA as such does not lead to an enrichment in long-distance cross-links. However, it could be that mass-corrected precursors tend to have a higher proportion of long-distance links. We therefore split the Xi-MPA results into five groups corresponding to the monoisotopic mass change (0, −1, −2, −3, −4 Da) and looked at their match to crystal structures. If a link originated from PSMs with different mass corrections, all of those were considered. We conducted a ‘nonparametric ANOVA’ (Kruskal–Wallis test) to detect any significant changes in the distance distributions of Xi-MPA identifications with different shifts and decoy distribution. However, we fail to reject the null hypothesis at the predetermined significance level of *α* = *0.05* (p-value: 0.13), indicating that the distance distributions for all subsets are similar. This matches the visual inspection of distance distributions (Figure 3B). Furthermore, all individual distance distributions were significantly smaller than the derived reference distribution (one-sided Wilcoxon test, see **Table S5**). In conclusion, we do not see any evidence of in-search monoisotopic mass assignment leading to increased conflicts with crystal structures. We then evaluated the effect of in-search monoisotopic mass assignment on PSM quality as assessed by the search score. First, we compared the scores of PSMs with a mass shift (Xi-MPA identifications) to the scores of the same spectrum without a mass shift (uncorrected data). While scores with shifted mass have a median of 6.7, the corresponding decoy distribution. (D) Score distribution of PSM matches of the ‘decoy mass search’. Identifications with a positive mass shift generally follow the decoy distribution (note that there are correct identifications with a positive mass shift, albeit few, see Figure 1B) while identifications with a negative shift resemble unshifted identifications. The scores of negative-shifted PSMs are significantly higher than those of positive-shifted PSMs (one-sided Wilcoxon test, p-value < 2.2e-16).

median score is 2.3 when using the uncorrected masses (Figure 3C). As one would expect from an increased search space, the scores of decoy hits also improve, albeit only marginally. We find that the score difference of target PSMs is significantly larger than of decoy PSMs (one-sided Wilcoxon test, p-value: < 2.2e-16). We then turned to a ‘decoy mass search’ for which we not only searched the range from 0 Da to −4 Da, but also +1 Da to +4 Da. Assuming the monoisotopic peak in the uncorrected data is rarely lighter than the true monoisotopic peak, the new identifications should score like decoy identifications. Indeed, the resulting score distributions for targets with a positive mass shift follow the decoy distribution (Figure 3D). In contrast, identifications with a negative shift are distributed like the identifications without mass shift. In conclusion, in-search monoisotopic mass change leads to significantly improved scores with a distribution that resembles that of precursors that did not see a mass change (0 Da). Importantly, these improvements are not random since an equally large search space increase (+1 Da to +4 Da) results in a completely different score distribution that resembles the decoy distribution but not the distribution of identifications without a mass shift.

**Figure 3.**
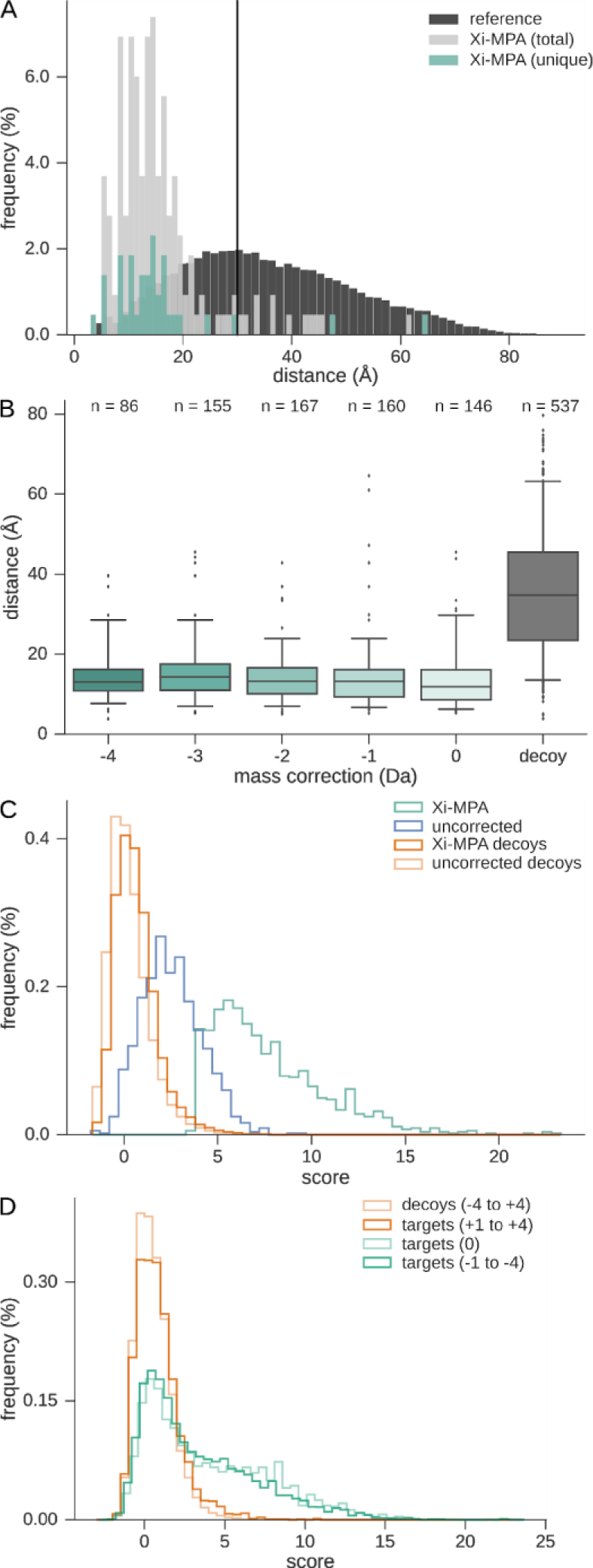
Matches with and without in-search mass shift show similar quality metrics. (A) Evaluation of Xi-MPA derived links on crystal structures (dataset 2). Distances between alpha carbon atoms of identified cross-linked residues in the crystal structure of the proteins are shown in light grey while a reference distribution of all possible pairwise C-alpha distances of cross-linkable residues is shown in dark grey. 30 Å is set as a limit, above which links are defined as long distance. (B) Distance distribution of identifications with different mass corrections. There was no significant difference between the different mass shifts, while all had a significant difference to the decoy distribution. (C) PSM scores of spectra identified with a mass shift are significantly higher than the corresponding score in uncorrected data. Shown are the score distributions of uncorrected and Xi-MPA results, as well as the corresponding decoy distribution. (D) Score distribution of PSM matches of the ‘decoy mass search’. Identifications with a positive mass shift generally follow the decoy distribution (note that there are correct identifications with a positive mass shift, albeit few, see Figure 1B) while identifications with a negative shift resemble unshifted identifications. The scores of negative-shifted PSMs are significantly higher than those of positive-shifted PSMs (one-sided Wilcoxon test, p-value < 2.2e-16).

### Heavy and low intensity peptides are corrected more frequently

One would especially expect to observe shifted mass assignment for peptides of high mass and low abundance. For large peptides (approximately >2000 Da), the monoisotopic peak will not be the most intense peak in the isotope cluster. If the peptide is of low abundance, the monoisotopic peak may be of too low intensity to be detected. We therefore analyzed the monoisotopic peak assignment in Xi-MPA regarding the precursor mass and intensity. Indeed, precursors with higher masses are more often corrected to lighter monoisotopic peaks (Figure 4A). While the median precursor mass for uncorrected matches is 2952 Da, for matches corrected by −2 Da it is already 4062 Da and for −4 Da it is 4684 Da. Of the identifications with a mass above 3000 Da, 88% were identified with a lighter mass. For precursors lighter than 3000 Da, the proportion was 42%. Like mass dependency, there is a trend towards larger correction ranges for lower intensity peptides (**Figure S5**).

**Figure 4.**
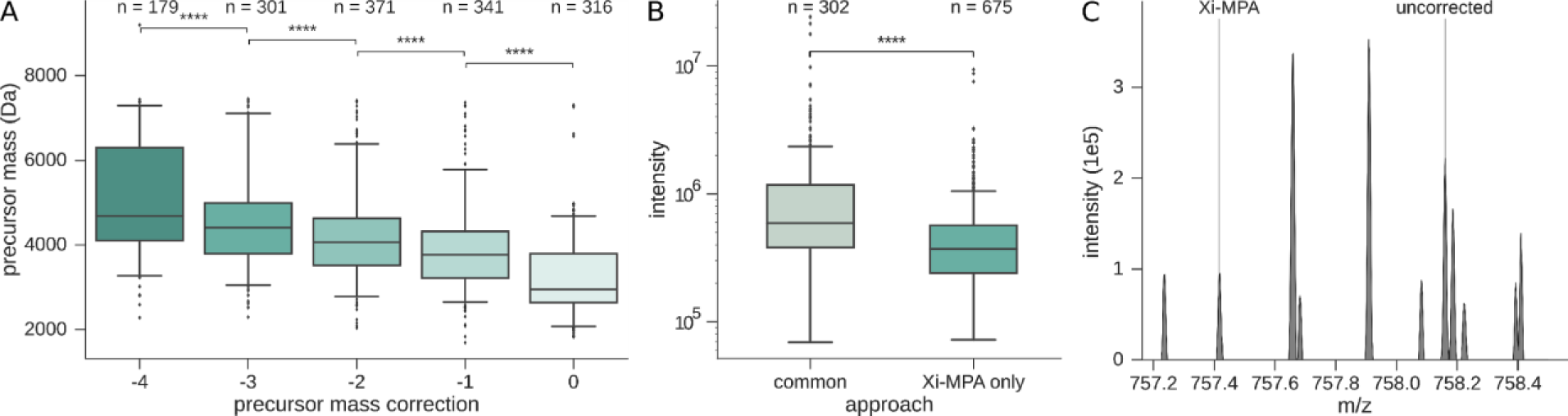
Correction is dependent of precursor mass and intensity. (A) Box plot of the precursor mass and monoisotopic mass correction of identified PSMs after Xi-MPA. PSMs with higher mass more often require monoisotopic mass correction to lighter masses. Whiskers show the 5 and 95% quantiles of the data. Asterisks denote the significance calculated by a one tailed t-test (****: p-value < 0.0001). (B) Precursors of cross-linked PSMs identified in all three approaches, MaxQuant-Xi, OpenMS-Xi, and Xi-MPA (‘common’), are more intense than precursors of PSMs that are only identified in Xi-MPA. In other words, successful correction happens more often for abundant precursors, while Xi-MPA identifies precursors of lower intensity. (C) MS1 isotope cluster of a cross-linked peptide. The monoisotopic peak of m/z 758.16 (z = 4, m = 3028.6 Da) was falsely assigned during acquisition and not corrected in any preprocessing approach. Xi-MPA identifies a PSM for a precursor with a mass that is 3 Da lighter.

When evaluating the newly matched precursors of Xi-MPA, the advantage of not having to rely on MS1 identification is evident. Matches not made through any of the preprocessing methods are generally much less intense (Figure 4B) and larger (**Figure S6**) than matches that are common to all approaches.

Manual analysis of isotope clusters of corrected precursors from dataset 2 revealed many cases where the monoisotopic peak was present in the MS1 spectrum but was not recognized during acquisition. For some, this might be due to the peak being of low intensity and discarded as noise, or because of other interfering peaks (Figure 4C). However, there are also cases where the cluster is well resolved (Figure 1C). Without details of how the instrument software determines the monoisotopic peak, a full evaluation is difficult. For a complete list of precursor m/z for Xi-MPA identifications and corresponding m/z of uncorrected, MaxQuant and OpenMS data, see **Table S1-S3**.

Note that in many acquisition methods, the machine only fragments peaks where it can successfully identify a full isotope cluster. Therefore, there might be instances of cross-linked peptides not being fragmented because of insufficient isotopic cluster quality, leading to lost identifications.

## CONCLUSION

The size and low abundance of cross-linked peptides leads to frequent misassignment of the monoisotopic mass by instrument software, which in some instances even escapes correction by sophisticated correction approaches employed by MaxQuant and OpenMS. Considering multiple monoisotopic masses during search increases the number of cross-link PSMs 1.8 to 4.2-fold, without compromising search accuracy as judged by multiple assessment strategies including comparison of the gains against solved protein structures. The problem of wrongly assigned monoisotopic peaks will have an impact on most cross-link search engines since these all rely in some part on the precursor mass. The extent of the misassignment will however be sample and software-dependent. Even with improved acquisition or correction software, there will remain instances where the monoisotopic peak cannot be determined correctly before searching due to low intensity. Our search-assisted monoisotopic peak assignment provides a general solution to this problem by relying on MS2 identification in addition to precursor information.

## ASSOCIATED CONTENT

### Supporting Information

Precursor m/z of different processing methods (XLS).

MS1 based mass range reduction, OpenMS preprocessing workflow, performance of Xi-MPA on additional data, dependency of correction on precursor intensity (PDF).

### Notes

The Authors declare no competing financial interest.

## ACKNOWLEDGMENT

We thank Dr Francis O’Reilly for comments and helpful discussions. This work was supported by the Einstein Foundation, the DFG [RA 2365/4-1], and the Wellcome Trust through a Senior Research Fellowship to JR [103139] and a multi-user equipment grant [108504]. The Wellcome Centre for Cell Biology is supported by core funding from the Wellcome Trust [203149].

## Table of Contents graphic

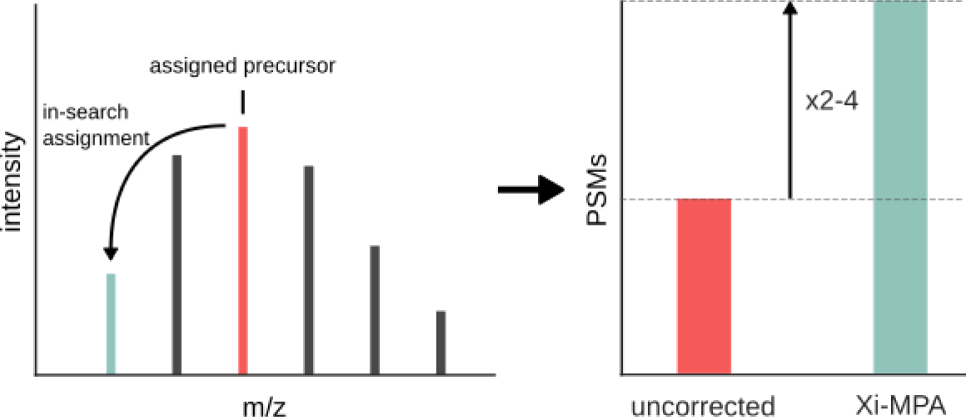

